# A Bottom-Up Platform for Quantitative Single-Particle Tracking Through Bacterial Biofilm Mimics

**DOI:** 10.64898/2026.07.02.736016

**Authors:** J.W. Shepherd, J.A.L. Howard

**Affiliations:** School of Physics, Engineering and Technology, University of York, York, UK

**Keywords:** Bacterial Biofilms, Biofilm Mimics, DNA Hydrogel, eDNA, Single Particle Tracking, Quantum Dots, Molecular Transport, SlimVar microscopy

## Abstract

Chronic infections persist in large part thanks to protection that biofilms afford their bacterial creators. The extracellular polymeric substance of biofilms is a hydrated matrix of DNA, polysaccharides, and structural proteins, amongst other components, through which nutrients, signalling molecules, and antimicrobial agents must diffuse to reach the bacteria within. Quantitative measurement of transport on the nanoscale within *in vivo* biofilms remains challenging due to optical heterogeneity, autofluorescence, active remodelling of biofilms and the ambiguity in trajectory reconstruction during single-particle tracking (SPT). Here, we present a methodological framework for measuring molecular transport in defined minimal extracellular matrix models using quantum dots as fluorescent nanoscale probes imaged with high-speed SlimVar microscopy.

To establish conditions in which high-diffusivity particle trajectories can be reliably reconstructed, upper limits to quantum dot concentrations were estimated from Brownian motion. The 99th-percentile inter-frame jump distance was estimated from the three-dimensional Brownian jump distance distribution and used to define a target average nearest neighbour distance, and therefore a per-particle volume, used for calculating a concentration which minimises the probability of trajectory collision during data acquisition. Quantum dot movement was imaged at sub-millisecond frame rates and diffusion coefficients were calculated in a 20% glycerol control and in DNA nanostar hydrogels modelling minimal extracellular matrix scaffolds assembled at 250 μM and 500 μM. Median diffusion coefficients decreased from 94.9 um^2^·s^-1^ in glycerol to 15.9 um^2^·s^-1^ and 8.3 um^2^·s^-1^in the 250 μM and 500 μM hydrogels, respectively.

More broadly, this work establishes a workflow for quantitative SPT in minimal biofilm models. Rather than attempting to reproduce the full biological complexity of native biofilms, this approach provides the basis of a modular experimental framework in which individual extracellular matrix components can be incorporated sequentially and their effects on molecular transport quantified.

## Introduction

The efficacy of antimicrobials relies on both their mechanism of action and their ability to reach their intended target; in general, they must traverse the extracellular environment before they can act. Transport dynamics therefore represents the first physical process governing therapeutic effectiveness. Theoretical analyses first showed the importance of molecular diffusion, sorption and interaction in determining antimicrobial progress (1). This becomes additionally important in biofilms because of the extracellular polymeric substance (EPS) surrounding bacterial cells, forming a physicochemical barrier between the environment and the bacterial community, through which every nutrient, metabolite, and antimicrobial must first diffuse before interacting with resident cells (2–6).

Biofilms have been shown to contribute to persistent infections in chronic wounds and cystic fibrosis, infective endocarditis, and infections associated with implanted medical devices (2, 4, 7). The increased resilience of pathogens within biofilms arises from multiple interacting biological processes, including altered bacterial metabolism, phenotypic heterogeneity, stress responses, persister cell formation, and the blockage of antimicrobial penetration (8–10). Although antibiotic diffusion dynamics alone do not explain increased antimicrobial resistance, every antimicrobial must traverse the EPS. Transport therefore plays a major role in the physical processes governing biofilm function. Imaging this transport through biofilms or *in vitro* models provides an experimentally accessible means of investigating biofilm matrix architecture.

The EPS is now understood to be a highly organised biological system consisting of extracellular DNA (eDNA), polysaccharides, and structural proteins, among other components. The relative abundance of each of these components can vary with species, biofilm developmental stage, and environmental conditions (5, 6, 11, 12).

These components collectively determine pore structure, hydration, the electrostatic environment, and the distribution of transient molecular interactions experienced by diffusing species. Because of this, transport through biofilms cannot be understood solely in terms of viscosity or geometric obstruction, especially as the biofilm architecture itself evolves during biofilm development.

eDNA is one of the most significant structural components of the EPS, as evidenced by DNaseI treatment preventing biofilm formation in *Pseudomonas aeruginosa*. (13). eDNA has subsequently been shown to contribute to biofilm cohesion, self-organisation and mechanical stability (14, 15). eDNA is also able to form electrostatic complexes with cationic polysaccharides such as Pel (16), and can directly prevent the penetration of antibiotics including vancomycin (10).

Considerable effort has been devoted to measuring transport within biofilms. Fluorescence recovery after photobleaching (FRAP), fluorescence correlation spectroscopy (FCS), penetration assays and microrheological approaches have all provided important insights into the mobility of solutes and tracer particles within extracellular matrices (17). These ensemble measurements have demonstrated that diffusion within biofilms is frequently slower than in bulk solution and may vary substantially with matrix composition (18). However, ensemble techniques necessarily average over spatial heterogeneity. Single-particle tracking (SPT) instead follows the trajectories of individual probe particles, enabling the direct resolution of local diffusion coefficients, anomalous transport, transient confinement, and population heterogeneity (19–23). Notably, one investigation using SPT in native bacterial biofilms revealed pronounced spatial heterogeneity (24).

Because biofilms are biologically rich systems, they remain difficult environments in which to establish causal relationships between matrix composition and transport.

Biofilms are dynamic, heterogeneous and continually altered by their resident microorganisms. The composition of the biofilm matrix continually changes throughout biofilm development and varies between bacterial species and environmental conditions (7, 25). As a result, experiments performed in living biofilms invariably assay many simultaneously varying physical and biological parameters. Changes in transport may arise from altered polymer composition, crosslink density, bacterial physiology, metabolic state or active matrix remodelling, making interpretation of *in vivo* biofilm experiments challenging.

Mechanistic biophysical understanding of complex systems often emerges from experimentally tractable model systems in which individual variables can be isolated and investigated quantitatively. For example, artificial lipid bilayers (26, 27), *in* vitro reconstituted cytoskeletal networks (28), while synthetic extracellular matrices have become indispensable tools for dissecting the influence of mechanical and biochemical cues on cell behaviour (29–31).

Applying this approach to bacterial biofilms requires constructing experimentally tractable extracellular matrices in which individual structural components can be introduced sequentially while their influence on transport is quantified. This is an idea already being used in artificial biofilm systems that employ defined polymeric materials to investigate selected matrix functions (32–34).

eDNA provides an attractive foundation for this kind of minimal model because of its key role in EPS structure and its exceptional experimental programmability. DNA hydrogels can be assembled from well-defined molecular building blocks whose topology, connectivity and mechanical properties can be tuned with molecular precision (35–38), and therefore represent a matrix model into which EPS components including polysaccharides, DNA-binding proteins, multivalent ions and, ultimately, living bacterial communities and antibiotics can be introduced with total control of physicochemical parameters like ionic strength and pH. Hydrogels are also an attractive system because of their practicality. They are cheap and quick to make, can be imaged with fluorescence microscopy, and do not need special health and safety handling as a full bacterial model would.

Here we establish a quantitative experimental platform for single-particle tracking through DNA hydrogel matrices. We use programmable DNA nanostar hydrogels to provide a minimal extracellular scaffold mimic, employing high-speed SlimVar microscopy(39, 40) to enable rapid imaging of diffusing quantum dots within optically thick specimens, with trajectories reconstructed and physical parameters extracted using bespoke analysis pipelines (41). To ensure the accurate linking of quantum dot trajectories, probe concentrations were selected using an analytical framework based on Brownian displacement statistics and the effective three-dimensional imaging volume, providing a conservative criterion that minimises trajectory ambiguity during rapid diffusion. Together, these methods establish a bottom-up platform for quantitatively linking matrix architecture to molecular transport and provide a foundation for constructing increasingly sophisticated biofilm mimics.

## Methods

### Theoretical Bounding of Quantum Dot Concentrations

To estimate the upper-bound for working concentration of quantum dots, we estimated the relevant length scales of diffusion in one time step. From Brownian motion, we know that the expected jump distance is given by a normal distribution with mean 0 and variance

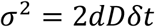

where d is the number of dimensions of diffusion, here 3, D is the diffusion coefficient, and δ*t*is the time between samples, here the frame-to-frame time on the camera.

Equivalently, the standard deviation of the jump distance is (41, 42)

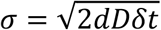

To be reasonably certain that successive spot localisations represent the same molecule, we estimate the upper bound of concentration so that on average the distance between particles is a 99th-percentile jump, which corresponds to 3σ. Clearly here the relevant dimension is *z*, given that we have a depth of field of approximately 500 nm (39), and *x* and *y* side lengths 256 and 66 pixels (or 16.64 and 4.29 microns) respectively. We therefore consider the minimal volume per particle to be one *xy* field of view, with a height calculated as 3σ + the depth of field of the microscope + 3σ-i.e. the appropriate distance both above and below the depth of field, while the height of the depth of field is also accounted for (See Figure 2a), so that if tesselating the schematic in Figure 2a there would be 6σdistance between particles vertically. As a result, our estimate for concentration upper bound is linked not only to the camera frame rate but the diffusive properties of the particle being tracked. To avoid circular reasoning, i.e. measuring diffusion coefficient, using that in the above calculation, and working out the required concentration to measure the diffusion coefficient, we calculated a range of estimated concentration upper bounds using a range of diffusion coefficients up to 200 µm^2^·s^-1^, which is well beyond what we would expect to measure in this system.

### DNA Hydrogel and sample preparation

DNA hydrogels were prepared using the nanostars described in (38). Briefly, three oligonucleotides: nanostar1 – 5’ CTGGATCCGCGGAAGCTTAAAACGGAATTCGCATGGATCCCCACGATCG nanostar2 – 5’ GGGGATCCATGCGAATTCCGAACTGAATTCCCTGGGATCCCGACGATCG nanostar3 – 5’ CGGGATCCCAGGGAATTCAGAATTAAGCTTCCGCGGATCCAGACGATCG containing 20 bases of complementarity to each other, either side of a flexible AA pair, and a terminal 6 base pair sticky end, were resuspended in Milli-Q water before being mixed in equimolar amounts. The oligonucleotide mix was then lyophilised in a vacuum drier before being resuspended in the necessary volume of annealing buffer (40 mM Tris (pH 8.0), 40 mM sodium acetate, 1 mM EDTA and 150 mM NaCl) to make final concentrations of 555 or 278 µM. The mixes were then annealed by heating to 90 °C in a covered heat block for 5 minutes, then allowed to cool slowly to room temperature overnight.

Slides and coverslips were prepared by plasma cleaning using atmospheric plasma for 1 minute, and flow cells were created by sticking coverslips to the slide using two pieces of double-sided tape separated by ∼5 mm, giving a chamber height of 120 μm. Quantum dots (PlasmaChem PL-QDN-590) were resuspended in 20 mM Tris pH 8.0 and diluted to 50 pM in annealing buffer. Quantum dots and nanostar hydrogels were then mixed in a 1:9 volumetric ratio, giving final concentrations of 5 pM quantum dots and either 500 or 250 µM nanostars as hydrogels. The hydrogel mix was then flowed into the flow cells using capillary action and the chambers sealed, ready for imaging.

The CdTe quantum dots had a manufacturer-specified inorganic core diameter of 3.3 nm and a hydrophilic, carboxyl-terminated (–COOH) surface. Hydrodynamic size was measured by dynamic light scattering (DLS) at 21 °C in the annealing buffer used for the SPT experiments, with buffer-only measurements used as controls. A quantum dot-associated small-particle population was identified from peaks in the intensity-weighted size distribution that were present following addition of quantum dots but absent from the buffer-only controls. The mean hydrodynamic diameter was 4.55 ± 0.72 nm (mean ± SE, n = 3). The corresponding free-solution diffusion coefficient in annealing buffer, estimated using the Stokes–Einstein relation and the viscosity of water at 21 °C, was 97 ± 15 µm^2^·s^−1^.

### SlimVar Microscopy

Quantum dots were imaged using a bespoke single-molecule super-resolved microscope capable of high-speed SlimVar microscopy with sub-millisecond temporal resolution when imaging bright inorganic dyes and quantum dots (39, 43), Figure 1a illustrates how the tilted angle of the laser excitation reduces background fluorescence emission signal. As shown schematically in Figure 1b, a 50 mW 532 nm laser (Coherent OBIS LS 532 nm 50 mW Laser System) is relayed through a 1:1 Keplerian telescope consisting of L_1_ and L_2_ (both f = 300 mm achromatic doublet AC254-300-A-ML, ThorLabs Inc.) with L_2_ being laterally translatable such that the beam can be steered from epifluorescence to a variable angle for SlimVar imaging. The laser is then focussed onto the back focal plane of the objective lens (Nikon CFI Apo TIRF 100X NA1.49 Oil) by L_3_ (f = 300 mm LB1779-A-ML lens, ThorLabs Inc.) via the dichroic mirror M_1_ (FF562-Di03, Semrock). Fluorescence emission is then collected via the objective lens and transmitted via M_2_ to the tube lens L_4_ (f=200 mm) within the microscope body (Nikon Eclipse Ti). A further 1:1 Keplerian telescope (L_5 &_ L_6_ both achromatic doublet lenses AC254-150-A with f = 150 mm, ThorLabs Inc.) transmits the image to a TeleDyne Photometrics Kinetix 22 sCMOS camera via emission filter F_1_ (ET585/65 M, Chroma). This produces a magnification of 100X and a resulting imaging pixel size of 65 nm. The Kinetix 22 camera has a fast row read time of ∼4 μs which allows for the sub-millisecond imaging of fluorescent molecules with a long enough exposure time for them to be visualised above the background signal. Images were acquired with a field of view consisting of 256 columns (approximately the size of the full width at half maximum of the laser excitation profile) and 66 rows, in 12-bit sensitivity mode and PM Cam - All Rows expose out, giving a frame-to-frame time of 0.854 ms. Quantum dots were tracked using the PySTACHIO software package (41). Tracks were generated when quantum dots could be linked over at least two frames, whilst apparent diffusion coefficients were subsequently estimated using the covariance-based estimator (CVE) of Vestergaard et al.(44), for which at least three consecutive localisations (two displacements) were required. A missing localisation terminated a trajectory and so quantum dot blinking could fragment trajectories or cause tracks to fall below the CVE inclusion threshold because we do not perform any dark interval linking.

**Figure 1:**
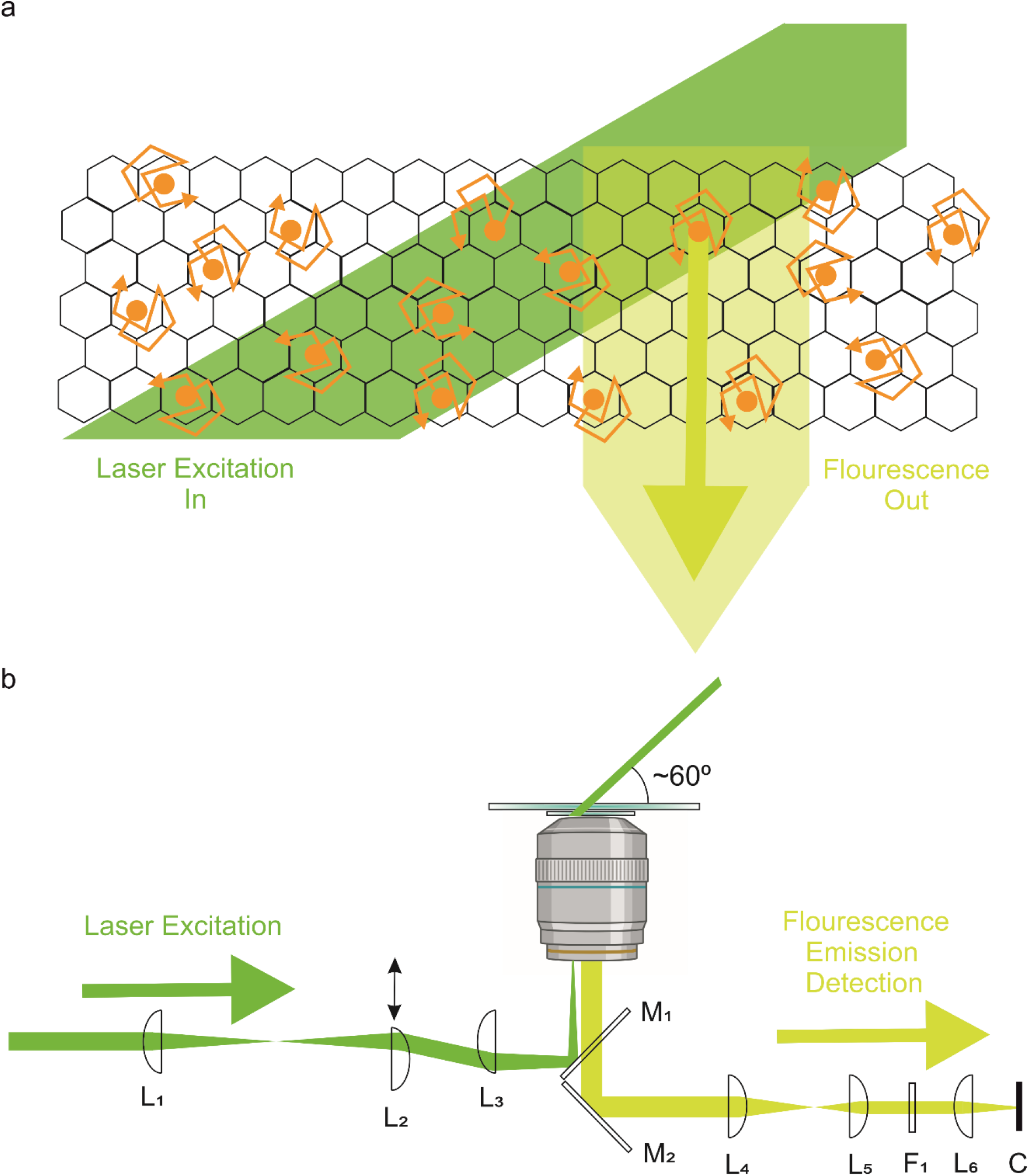
**a)** schematic representation of SlimVar microscopy: the tilted angle of the laser excitation means that fewer fluorescent molecules are excited in the region where light is collected by the objective, resulting in a much lower background signal and, as such, a higher signal-to-noise ratio. **b)** the optical path taken by the laser and subsequent fluorescent emission

## Results

### The upper bound of tracked particle concentration depends on the diffusion coefficient and frame-to-frame time

We find that as expected, there is a strong dependence of upper-bound concentration estimate on both diffusion coefficient and camera frame-to-frame time (see Figure 2b).

**Figure 2:**
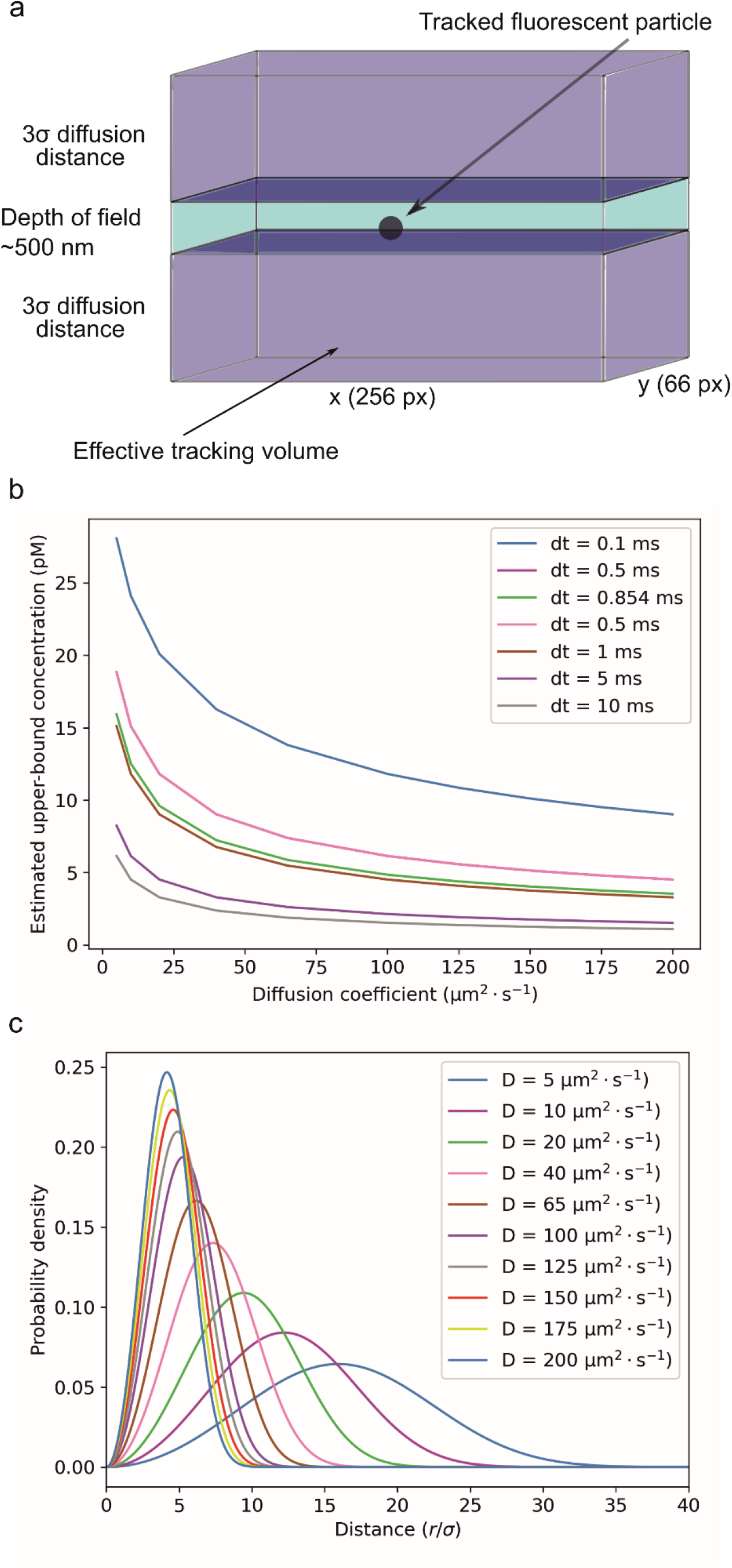
**a)** Schematic diagram of the spatial geometry used for estimating the required per-particle volume. In the centre of the light region a particle is diffusing inside a volume with height equal to the depth of field of the camera, here 500 nm (39). The dark regions above and below the light field of view indicate the additional volume to maintain average spatial separation giving Z_eff_=Z_DOF_+6σ. **b)** Estimated upper bound for concentration as a function of diffusion coefficient for a range of camera exposure times. We find that in the extreme case of 200 µm^2^·s^-1^, the estimated concentration is around 5 pM for sub-ms tracking. **c)** Probability density distribution for nearest-neighbour distance for various diffusion coefficients at dt = 0.854 ms. We find that all modal values are above 4 sigma.

For a camera frame-to-frame time of 0.1 ms, we estimate that even at 200 µm^2^·s^-1^, a concentration above 10 pM may be suitable, while for a camera rate of 10 ms, less than 1 pM is acceptable. Practically speaking, these estimates need to be tensioned against the need to collect data – a concentration of 0.001 pM is experimentally infeasible because it would be impossible to image for long enough to build up a dataset. In the relevant time range for this work, frame-to-frame time of 0.854 ms, we find an estimate of 4.8 pM for a 100 µm^2^·s^-1^ diffusion coefficient. Experimentally, we found that a concentration of 5 pM provided confidence in the data collection, as well as a reasonable number of single particle tracks for analysis. We note that our calculations here represent a very conservative estimate upper bound on the usable working concentration: in Figure 2c the probability density function of nearest neighbour distance (40, 45) is plotted for our chosen frame-to-frame time of 0.854 ms and various diffusion coefficients. We see that the modal nearest neighbour separation is well above 3σ in all cases.

### Matrix density modulates single-particle transport

Single-particle trajectories were analysed to obtain apparent diffusion coefficients (D_app_) for individual quantum dots in 20% glycerol and DNA nanostar hydrogels. The resulting distributions were strongly right-skewed when expressed in linear units, with a small number of rapidly diffusing particles extending the distribution towards several hundred µm^2^·s^-1^. Logarithmic transformation substantially reduced this skewness, yielding broad but approximately unimodal distributions suitable for quantitative comparison (Figure 3a). Relative to 20% glycerol, both DNA nanostar hydrogels shifted the median D_app_ towards lower values. The median D_app_ decreased from approximately 94.9 µm^2^·s^-1^ (IQR: 41.0 – 181.2 µm^2^·s^−1^) in 20% glycerol to 15.9 µm^2^·s^-1^ (IQR: 5.7 – 49.5 µm^2^·s^−1^) in the 250 µM nanostar hydrogel and 8.3 µm^2^·s^-1^ (IQR: 2.7 – 33.0 µm^2^·s^−1^) in the 500 µM hydrogel (Figure 3a), with all distributions having P-values < 0.0001 when compared using Holm-corrected Dunn’s post hoc test . This corresponds to approximately a six-fold reduction in the median D_app_ at 250 µM and an approximately eleven-fold reduction at 500 µM relative to 20% glycerol. Although diffusion slowed substantially in the DNA hydrogels, the distributions remained broad, spanning more than an order of magnitude in D_app_.

**Figure 3.**
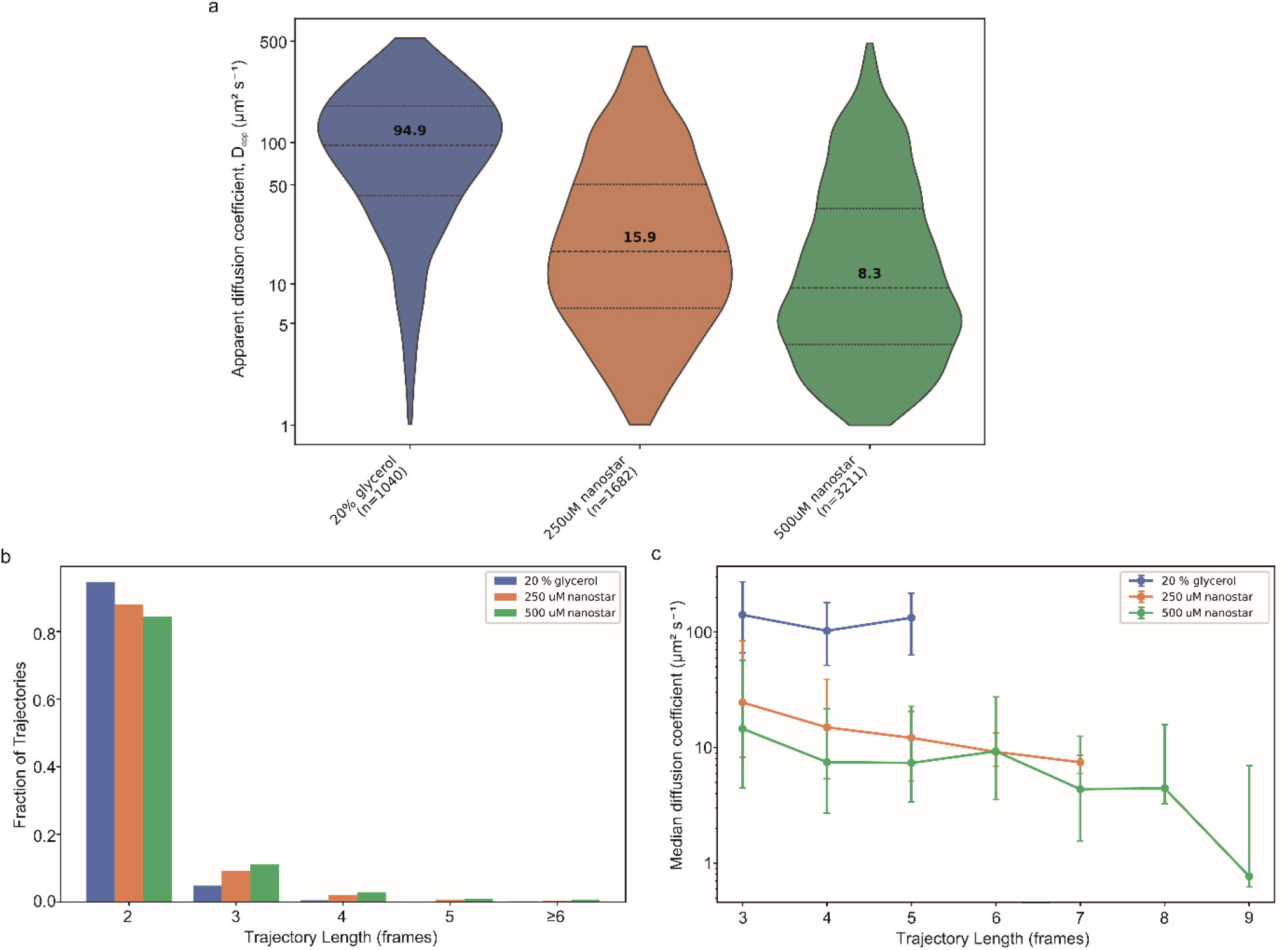
Effect of matrix density and trajectory length on apparent quantum-dot diffusion. **a)** Violin plots of CVE-derived apparent diffusion coefficients (D_app_) in 20% glycerol and 250 µM and 500 µM DNA nanostar hydrogels. Dashed horizontal lines indicate the median and dotted lines the interquartile range. Median values are annotated within each distribution. Distributions were compared using Dunn’s Post Hoc Test, and Holm corrected to give P-values of 2.1 x 10^-118^ (glycerol v 250 µM DNA nanostars), 1.7 x 10^-257^ (glycerol v 500 µM DNA nanostars) and 6.4 x 10^-25^ (250 v 500 µM DNA nanostars). **b)** Trajectory-length distributions for all reconstructed trajectories, shown as the fraction of trajectories in each condition. **c)** Median CVE-derived D_app_ as a function of trajectory length for trajectories meeting the CVE inclusion criterion of length at least 3 frames. Error bars show the interquartile range.

This indicates that individual particles experience markedly different transport environments despite being measured within nominally identical samples. Increasing nanostar concentration shifted the entire distribution towards lower D_app_ without producing a distinct immobile population, suggesting that increasing matrix density progressively restricts molecular transport rather than introducing a discrete trapping state.

We additionally examined trajectory length before CVE filtering (Figure 3b). All three conditions were dominated by short reconstructed trajectories, with the 20% glycerol condition showing the greatest enrichment of two-frame tracks as expected for the system with the highest diffusion coefficient where the quantum dots rapidly diffuse out of the imaged depth of field. Two-frame trajectories contribute to the track-length distribution but are not included in the CVE analysis, which requires at least three consecutive localisations.

To test whether D_app_ itself varied systematically with the length of accepted trajectories, median D_app_ was plotted as a function of trajectory length (Figure 3c). The ordering between conditions was retained across the trajectory lengths tested, with glycerol showing the highest D_app_, followed by 250 µM and then 500 µM nanostar hydrogels. The consistent reduction in D_app_ between these hydrogels within matched trajectory-length classes indicates that the concentration-dependent reduction in apparent mobility is not produced solely by the minimum-track-length criterion.

## Discussion

This study establishes a reductionist experimental platform for investigating molecular transport through defined extracellular matrices using single-particle tracking.

Combining DNA nanostar hydrogels with high-speed SlimVar microscopy, we show that increasing matrix density produces a systematic reduction in the mobility of individual quantum dots while preserving the heterogeneity of transport. We also introduce a quantitative framework for selecting probe concentrations based on Brownian statistics, allowing single-particle measurements to be performed while minimising the chance of mis-linking trajectories. Together, this provides a robust experimental system in which the relationship between extracellular matrix architecture and molecular transport can be investigated under precisely controlled conditions.

The measurements in 20% glycerol also define an important practical limitation of the present SPT implementation. The DLS-derived hydrodynamic diameter predicts a free-solution diffusion coefficient of approximately 97 ± 15 µm^2^·s^−1^ in annealing buffer at 21 °C. Direct buffer-only SPT at this diffusivity would be expected to be strongly affected by motion blur, finite axial residence and trajectory-linking ambiguity. The 20% glycerol condition therefore represents an experimentally accessible fast-diffusion/known-viscosity control rather than a zero-matrix baseline. Using the measured hydrodynamic diameter and the viscosity of 20% (v/v) glycerol at 21 °C (1.9292 mPa·s), the Stokes– Einstein relation predicts D ≈ 49 ± 8 µm^2^·s^−1^, whereas the measured median CVE-derived D_app_ was 94.9 µm^2^·s^−1^ (IQR: 40.95 – 181.2 µm^2^·s^−1^). Together with the predominance of very short glycerol trajectories, this discrepancy indicates that the fast-diffusion control approaches the practical limits of trajectory reconstruction. The absolute glycerol D_app_ should therefore not be interpreted as an unbiased estimate of free quantum dot diffusion. These limitations affect the interpretation of the absolute fast-control value, but the reduction in D_app_ between the 250 and 500 µM nanostar hydrogels persists within matched trajectory-length classes and therefore supports a matrix concentration-dependent restriction of transport.

An important advantage of this reductionist framework is that additional matrix components can be incorporated sequentially while maintaining experimental control. In *in vivo* biofilms, extracellular DNA rarely exists in isolation, instead interacting with a multitude of other components such as polysaccharides, proteins, multivalent ions and DNA-binding proteins that collectively determine matrix architecture. Among these, the bacterial nucleoid-associated protein Integration Host Factor (IHF) is emerging as an interesting DNA binding protein to investigate. Structural and single-molecule studies have demonstrated that IHF binds DNA through multiple bending and bridging modes with varying sequence specificity, generating a diverse range of higher-order DNA conformations (46). It has also been shown to substantially modify the emergent mechanics of entangled DNA networks by promoting their fluidisation and structural rearrangement (47). Under acidic conditions characteristic of mature biofilms, IHF can undergo a transition from predominantly DNA bending to extensive intermolecular DNA bridging increasing network connectivity (48). These observations suggest that proteins acting at the level of individual DNA molecules may produce measurable changes in the bulk physical properties of extracellular matrices, including molecular transport.

The experimental platform presented here provides an attractive route for testing the effect that IHF and similar proteins have on extracellular matrices in combination with other co-factors. By introducing purified DNA-binding proteins into a chemically defined matrix while monitoring single-particle transport, it should become possible to quantify directly how specific molecular interactions influence transport behaviour without the confounding effects of simultaneous biological adaptation. More broadly, the modular nature of DNA nanostar hydrogels provides a framework for progressively reconstructing increasingly sophisticated extracellular matrices. Additional matrix constituents,including exopolysaccharides, extracellular proteins, divalent cations and bacterial nucleoid-associated proteins, can be introduced individually or in combination while preserving quantitative experimental control. The ability to quantify molecular transport within programmable extracellular matrices will be a powerful tool for understanding diffusion in a wide range of biological soft materials and synthetic biomaterials.

## Acknowledgements

J.W.S. and J.A.L.H. would like to thank the Leake group at the University of York for useful scientific discussions and Prof. Leake especially for mentorship and for generous free usage of microscopical equipment.

